# Detecting spatiotemporal pattern of tuberculosis and the relationship between ecological environment and tuberculosis, a spatial panel data analysis in Guangxi, China

**DOI:** 10.1101/348169

**Authors:** Zhezhe Cui, Dingwen Lin, Virasakdi Chongsuvivatwong, Jinming Zhao, Mei Lin, Jing Ou, Jinghua Zhao

## Abstract

Guangxi is one of the provinces having the highest reported incidence of tuberculosis (TB) in China. However, spatial and temporal pattern and causation of the situation are still unclear. In order to detect the spatiotemporal pattern of TB and the association with ecological environment factors in Guangxi Zhuang autonomous region, China, We performed a spatiotemporal analysis with prediction using time series analysis, *Moran’s I* global and local spatial autocorrelation statistics, and space-time scan statistics, to detect temporal and spatial clusters. Spatial panel models were employed to identify the influence factors. The time series analysis shows that the number of reported cases peaked in spring and summer and decreased in autumn and winter with the annual reported incidence of 113.1/100,000 population. *Moran’s I* global statistics were greater than 0 (0.363 – 0.536) during the study period. The most significant hot spots were mainly located in the central part. The east part exhibited a low-low relation. By spacetime scanning, the clusters identified were similar to that of the local autocorrelation statistics, and were clustered toward the early of 2016. Duration of sunshine, per capita gross domestic product (PGDP), the recovery rate of TB and participation rate of new cooperative medical care insurance in rural areas had a significant negative association with TB. In conclusion, the reported incidence of TB in Guangxi remains high. The main cluster was located in the central part of Guangxi, a region where promoting the productivity, improving TB treatment pathway and strengthening environmental protective measures (increasing sunshine exposure) are urgently needed.

## Introduction

Tuberculosis (TB) caused by Mycobacterium tuberculosis (MTB) is a severe public health problem worldwide, especially in developing countries. The burden of TB in China is high. According to World Health Organization (WHO),^[1]^ the number of TB cases reported from China was the third highest globally in 2016, after India and Indonesia. Guangxi is an autonomous region located in southern China and is also regarded as a high TB region with about 50,000 new TB cases reported by NNDRS every year. It is ranked as the third highest TB burden province in China. The annual reported incidence is about 100 cases per 100,000 population.

Geospatial analytical methods are essential tools for increasing our understanding of public health problems. There is an increasing number of studies that have used geospatial analytical methods to analyze disease trends and to detect the association between the health issues and other factors.^[2–4]^ The TB situation appears complex and spatially heterogeneous in China.^[5–6]^ Previous studies were based on the provincial and city levels.^[7–8]^ Until now, few studies have analyzed the spatiotemporal dynamics of TB clusters at the county level in China.^[9]^ Some areas of Guangxi have a higher reported incidence of TB than other areas. Conversely, some areas have a lower level. In addition, diversity of ecological environment in Guangxi indicates that there are quite possibly have some independent variables influencing the TB transmission. Unfortunately, studies of TB clustering in Guangxi have rarely been reported in the past. The mode of TB transmission and driving factors are still unclear. For these reasons, we performed a spatiotemporal analysis of TB in Guangxi between 2010-2016 using time series analysis, spatial statistical analysis, and space-time scan statistics to detect high and low reported incidence areas. Spatial panel data model were employed to identify the environmental influence factors.

## Methods

### Data collection

Guangxi has a population of about 46 million. It covers an area of 236,700 km^2^ and has a 1,020 km border with Vietnam to the south-west. Guangxi is divided into 14 cities containing 113 counties and districts. Currently, Guangxi has three TB control models. The first one is a Center for Disease Control and Prevention (CDC) model covering 51 counties/districts which integrates TB patient diagnosis, treatment and management into one package. The second is the designated hospital model covering 39 counties/districts which separates the TB patient diagnosis, treatment into the local designated hospital and the patient management into CDC. The third one is a specialist hospital model covering two districts which integrates TB patient diagnosis, treatment and management into one package in a TB specialist hospital. We collected the TB cases from the National Notifiable Disease Reported System (NNDRS) and those reported by General Hospitals, TB designated hospitals, CDC, and specialist hospitals, and verified by the National TB Program (NTP).^[10]^ NNDRS is a multifaceted program that includes a surveillance system for infectious diseases and sharing of data. Public health officers use this information to monitor, control, and prevent the occurrence and spread of nationally notifiable infectious diseases and conditions. The NNDRS records information including name, age, gender, current address, result of sputum test and final diagnosis of TB patients. TB diagnostic routines include symptoms screening, image examination, sputum smear and culture. If a patient is suspected as high risk for Multiple Drug Resistant Tuberculosis (MDR-TB), species identification, drug sensitivity testing and Xpert-MTB/RIF assay will be performed. We excluded patients who were diagnosed with Non-Tuberculosis Mycobacterial (NTM) infection by species identification. Confirmed TB cases are treated in TB designated hospitals, CDC or specialist hospital following the Directly-Observed Treatment Strategy.

Overall 365,179 confirmed TB cases during 2010-2016 in Guangxi were identified in this study. We aggregated the TB cases by county and matched them by area code. Annual population data for each administrative district was obtained from the Sixth Nationwide Population Census database in 2011. Ecological environment data included altitude (ALT), annual rainfall (AR), duration of sunshine (DOS), average temperature (AT), average humidity (AH), forest cover (FC), trees in woodland forest (TWF), trees in sparse forest (TSF), scattered trees (ST), trees planted by the side of farm houses, roads, rivers and fields (TPS), sex ratio (SR), total gross domestic product (TGDP), per capita gross domestic product (PGDP), TB control fund (TBF), health fund (HF), number of hospitals (NOH), number of grass-root health facility (NGHF), number of hospital beds (NHB), number of doctors (NOD), number of other health worker (NOHW), recovery rate of TB (RRTB), mortality of TB (MTB), prevalence of HIV/AIDS (PHIV) and participation rate of new cooperative medical care insurance in rural areas (PRR) were obtained from the China Meteorological Data Sharing Service System, statistical yearbook and health resource database. After data collection, we construct a panel data frame with the cross-section of observations repeated over several time periods. We obtained vector map files from the Global Administrative area database *(GADM Inc, California, US)*.

### Statistical analysis

#### (a) Time series

The number of TB cases was aggregated by month for trend analysis at time dimension. We conducted the time series analysis at the provincial level retrospectively. The time series included 84 months starting from January 2010 and ending in December 2016 and was examined by *IBM SPSS* statistical data editor (version 19.0, Statistical Product and Service Solutions, Chicago, IL). We used the “forecasting” function to form the sequence chart, and selected the better-predicted model. The predicted model was obtained, together with model diagnostics, and predicted values for the early-warning of TB reporting.

#### (b) Spatial Autocorrelation Analysis

*Moran’s I* enables us to measure spatial autocorrelations^[11]^ and has been widely used in the spatial analysis of TBJ^[12–13]^ Spatial autocorrelation statistics include global spatial autocorrelation and local spatial autocorrelation. The global spatial autocorrelation estimates the overall degree of spatial autocorrelation. The local spatial autocorrelation is able to identify the location and types of clusters. We used *GeoDa 1.8.12 (Luc Anselin, University of IL Linois, Urbana-Champaign, US)* to analyze the spatial autocorrelation. We also used the Empirical Bayes (EB) adjustment to take into account variance instability of rates in the global and spatial autocorrelation.

A *Moran’s I* value greater than 0 indicates that the reported incidence of TB in neighboring districts exhibit spatial autocorrelation compared to the non-neighboring districts. A value of 0 means that the results may vary slightly due to random permutation. A value less than 0 indicates that there is no significant clustering.^[14]^ A *P* value less than 0.05 suggests that we can reject the null hypothesis and conclude that there are some spatial autocorrelations in the study area. In order to obtain more robust results, we increased the number of permutations to 999.The global *Moran’s I* statistic is also the mean of the spatial *Moran’s I* statistic.

*GeoDa* can output the significance map which shows the locations with significant local Moran I statistics in different shades, and a local indicator of spatial association (LISA) cluster map. The high-high (red districts in the cluster map) and low-low (blue districts in the cluster map) locations are typically regarded to be TB spatial clusters, while the high-low and low-high locations (negative local spatial autocorrelation) are termed spatial outliers. While outliers are single locations by definition, this is not the case for clusters.

#### (c) Space-time Scan Statistic

In order to test spatial and temporal interaction and evaluate the relative risk for each cluster, we calculated the space-time scan statistics developed by Martin Kulldorff using *SaTScan*™ v 7.0.2. For continuous scan statistics, *SaTScan* uses a continuous Poisson model. ^[15]^ The scanning window is an interval (in time), a circle or an ellipse (in space) or a cylinder with a circular or elliptical base. Multiple different window sizes are used. For each location and size of the scanning window, the window with the maximum likelihood is the most likely cluster, that is, the cluster least likely to be due to chance. ^[16]^ The alternative hypothesis is that there is an elevated risk within the window as compared to outside. The space-time scan window with maximum likelihood value is defined to be the most likely cluster and other significant windows are defined to be secondary clusters.^[17]^ *R* version 3.3.2 was used to display the threedimensional visualization of the scan window.

#### (d) Spatial panel data models

The models for spatial panel data were used to estimate the association between the ecological environment factors and reported incidence of TB, which could analyze data with spatial dependence, and also enable us to consider spatiotemporal heterogeneity. we introduced the Spatial Panel Data Models to test the various spatial panel data specifications. Spatial panel data models capture spatial interactions across spatial units and over time. We consider the implementation of maximum likelihood estimators in the context of fixed as well as random effects spatial panel data models:
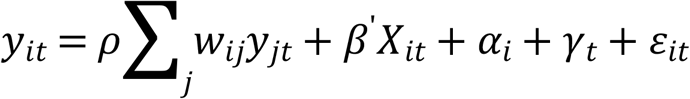

where *γ*_*t*_ is a year indicator. The spatial unit effect *α*_*i*_ captures unobserved time-invariant heterogeneity. It will decrease time-invariant bias from correlated effects. In this section, we used “splm” package of *R* to run spatial panel data Models. Before running the program, we created spatial weights matrix according to the neighboring relationship of each location. The splm object was a list of various elements including the estimated coefficients, the vector of residuals and fitted values. Baltagi, Song and Koh SLM1 marginal test was employed for random effects checking.

## Results

### Descriptive analysis of TB cases

Of the 365,179 active TB cases collected from NNDRS. The annual reported incidence of active TB was stable over the whole study period with obvious seasonal fluctuations. The mean annual reported incidence of active TB was 113.1/100,000 population (108.5 – 117.6). Annual active TB reported incidence shown downtrend (Chi-square for linear trend=159.76, *P*-value<0.001).

### Time series analysis

Figure 1 shows the monthly time series of active TB cases during 2010-2016 as well as the fitted curves and forecast curves for 2017 including Upper Control Limit (UCL) and Lower Control Limit (LCL) values. The type of forecast model was a simple seasonal model (Stationary *R*^2^=0.517, Normalized *BIC*=12.121). In general, TB cases peaked in spring and summer and decreased in autumn and winter. As shown in Table 1 the total number cases in 2016 were forecasted to exceed 49 thousand with a peak from March to July and the estimated annual incidence was 108.3 per 100,000 population.

**Figure 1.**
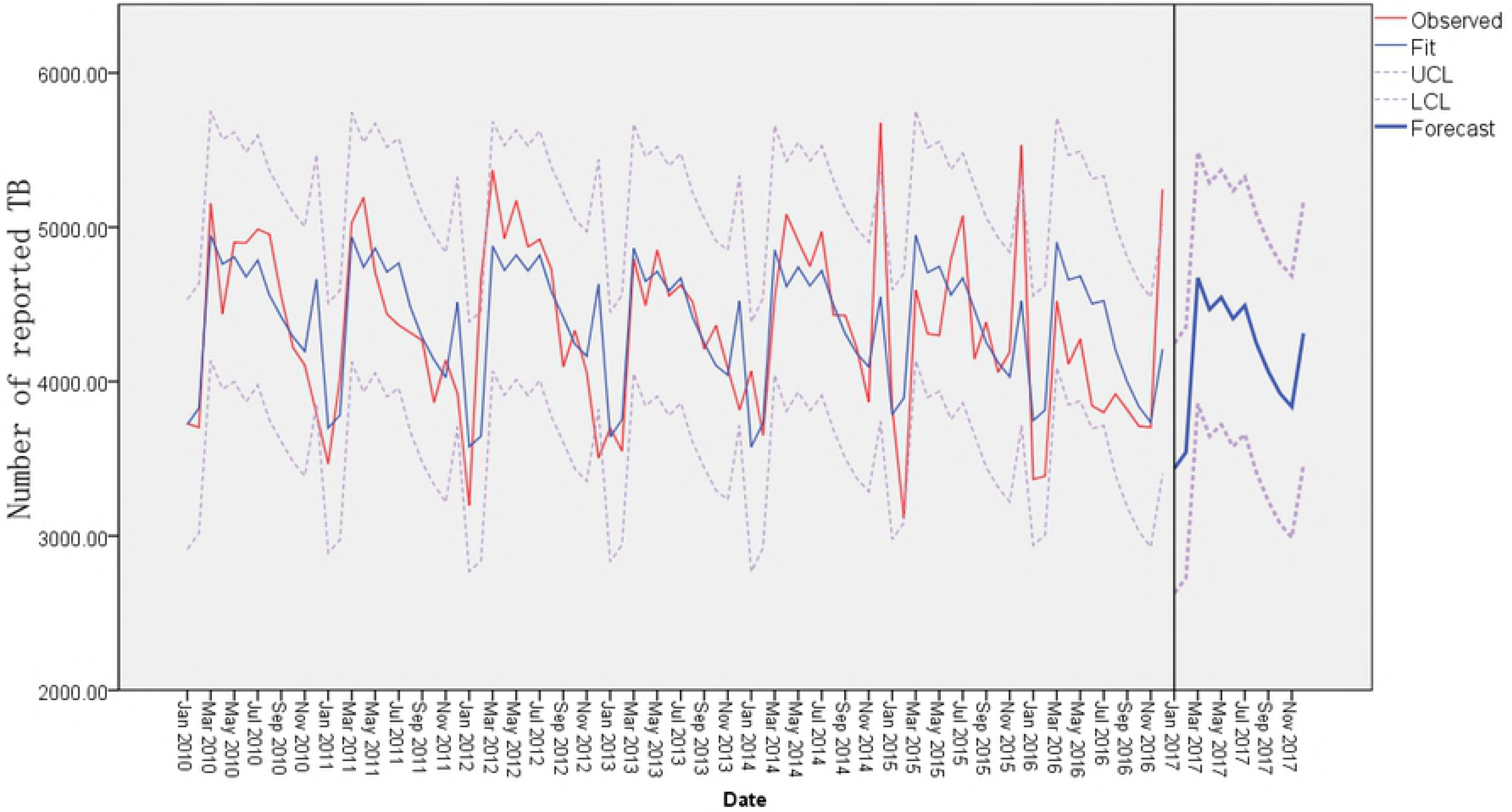
The monthly time series and forecast curve of TB cases in Guangxi, 2010-2017

**Table 1.** The forecast values for new TB cases in 2017

|  | Jan | Feb | Mar | Apr | May | Jun | Jul | Aug | Sep | Oct | Nov | Dec | Total |
| --- | --- | --- | --- | --- | --- | --- | --- | --- | --- | --- | --- | --- | --- |
| Forecast | 3437 | 3542 | 4672 | 4467 | 4545 | 4407 | 4492 | 4245 | 4066 | 3924 | 3836 | 4313 | 49946 |
| UCL | 4245 | 4355 | 5488 | 5287 | 5370 | 5236 | 5325 | 5081 | 4906 | 4768 | 4685 | 5165 | 59911 |
| LCL | 2628 | 2729 | 3855 | 3646 | 3721 | 3579 | 3660 | 3408 | 3225 | 3079 | 2988 | 3461 | 39980 |

### Spatial Autocorrelation Analysis

As shown in Table 2, all the *Moran’s I* statistics were greater than 0 (0.363 – 0.536). During the study period, the TB reported incidence of some neighboring districts exhibited spatial autocorrelation compared to the non-neighboring districts.

**Table 2.** Spatial Autocorrelation Analysis Results from 2010 to 2016

| year | Global spatial autocorrelation |  | Local spatial autocorrelation |  |  |  |  |
| --- | --- | --- | --- | --- | --- | --- | --- |
|  | *Moran's I Statistic* |  | high-high | low-low | high-low | low-high | no significance |
| 2010 | 0.536 |  | 19 | 21 | 2 | 1 | 69 |
| 2011 | 0.476 |  | 16 | 23 | 2 | 0 | 71 |
| 2012 | 0.512 |  | 15 | 25 | 1 | 0 | 71 |
| 2013 | 0.470 |  | 12 | 23 | 4 | 1 | 72 |
| 2014 | 0.520 |  | 12 | 22 | 2 | 2 | 74 |
| 2015 | 0.363 |  | 10 | 24 | 6 | 3 | 69 |

From the result of local spatial autocorrelation analysis with EB adjustment (Table 2, Figure 2), the most significant hot spots (high-high relation) were mainly located in the central part of Guangxi. Some neighboring districts such as Du’an, Yizhou, Xincheng and Xingbin consistently exhibited a high-high relation as the strongest hot spots in the period of observation. Counties or districts in the eastern of Guangxi exhibited a low-low relation (significant cold spots). The number of counties/districts exhibiting high-low and low-high were always less than 5. However, the pattern of TB clustering in some districts changed over the six-year study period. For example, two counties located in the northwest of Guangxi changed from non-significant to significant hot spots, while three counties in the southwest of Guangxi changed from significant hot spots to a non-significant area. The number of significant hot spots decreased from 19 in 2010 to 15 in 2016. In contrast, the number of counties/districts exhibiting significant cold spots in 2016 was more than that in 2010 (28 vs 21).

**Figure 2.**
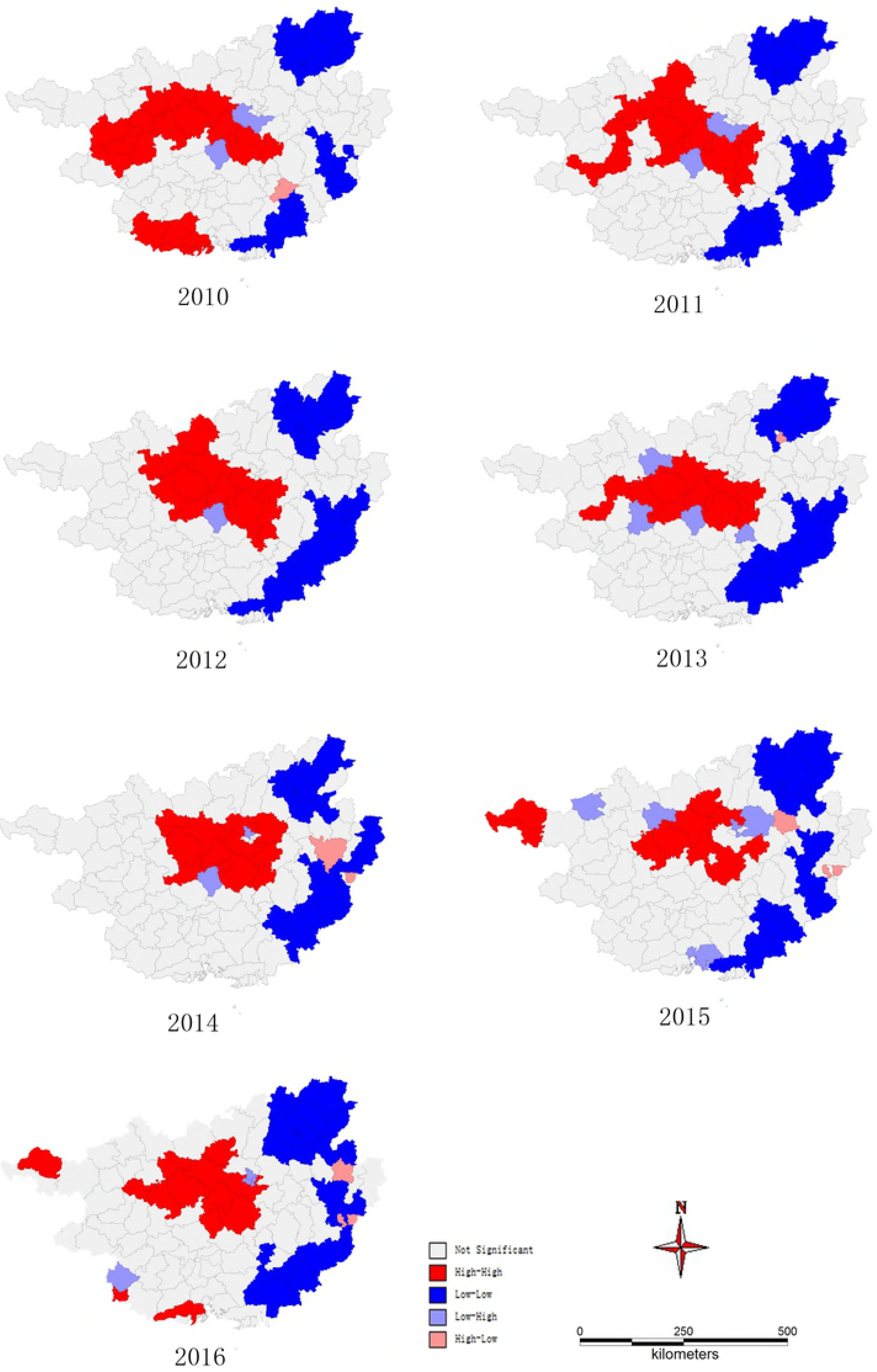
Cluster map for local spatial autocorrelation analysis in Guangxi, 2010-2016

### Space-time Scan Statistic

Table 3 shows space-time scan statistics in Guangxi from 2010 to 2016. There was one most likely cluster and three secondary clusters identified. Figure 3 shows that the most likely cluster and secondary cluster #3 were the high reported incidence rate districts (spatiotemporal hot spots) and included 17 counties/districts while the other two clusters were the low reported incidence rate districts (spatiotemporal spots) and included 34 counties/districts. As the most likely cluster, the center of the circle is Xincheng county with a radius of 77.3 km, and included nine counties. The cluster period persisted from February 2012 until July 2015. Residents of this region and during this time period had a 1.93 times higher risk of developing active TB compared to those living outside this region. The log likelihood ratio was also very high (4957.7). Another hot spot was located in the south of Guangxi near the Vietnamese border, and persisted from April 2014 until December 2015. The two cold spots were located in the eastern and northern parts of Guangxi, and persisted almost across the whole study period with a low relative risk (0.62 and 0.63).

**Table 3.**
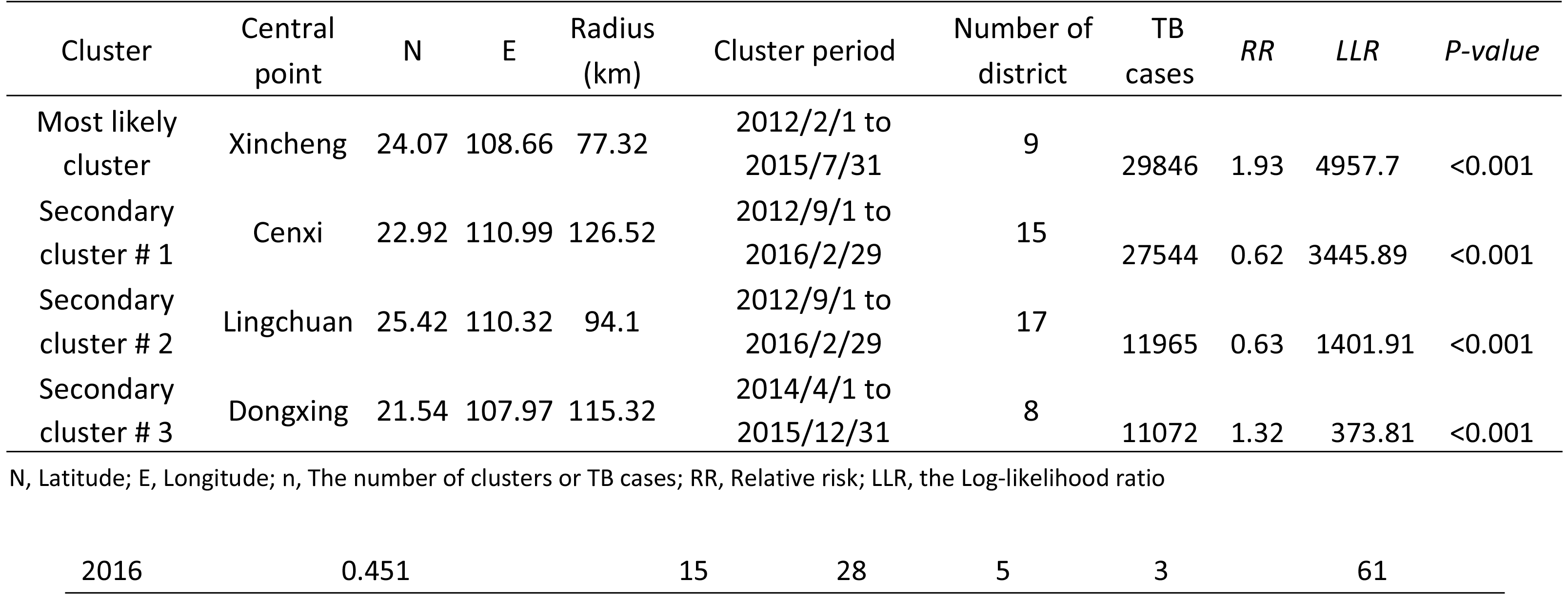
Space-time clusters of TB cases in Guangxi from 2010 to 2016

| Cluster | Central point | N | E | Radius (km) | Cluster period | Number of district | TB cases | RR | LLR | P-value |
| --- | --- | --- | --- | --- | --- | --- | --- | --- | --- | --- |
| Most likely cluster | Xincheng | 24.07 | 108.66 | 77.32 | 2012/2/1 to 2015/7/31 | 9 | 29846 | 1.93 | 4957.7 | <0.001 |
| Secondary cluster # 1 | Cenxi | 22.92 | 110.99 | 126.52 | 2012/9/1 to 2016/2/29 | 15 | 27544 | 0.62 | 3445.89 | <0.001 |
| Secondary cluster # 2 | Lingchuan | 25.42 | 110.32 | 94.1 | 2012/9/1 to 2016/2/29 | 17 | 11965 | 0.63 | 1401.91 | <0.001 |
| Secondary cluster # 3 | Dongxing | 21.54 | 107.97 | 115.32 | 2014/4/1 to 2015/12/31 | 8 | 11072 | 1.32 | 373.81 | <0.001 |

N, Latitude; E, Longitude; n, The number of clusters or TB cases; RR, Relative risk; LLR, the Log-likelihood ratio
|  |  |  |  |  |  |  |
| --- | --- | --- | --- | --- | --- | --- |
| 2016 | 0.451 | 15 | 28 | 5 | 3 | 61 |

**Figure 3.**
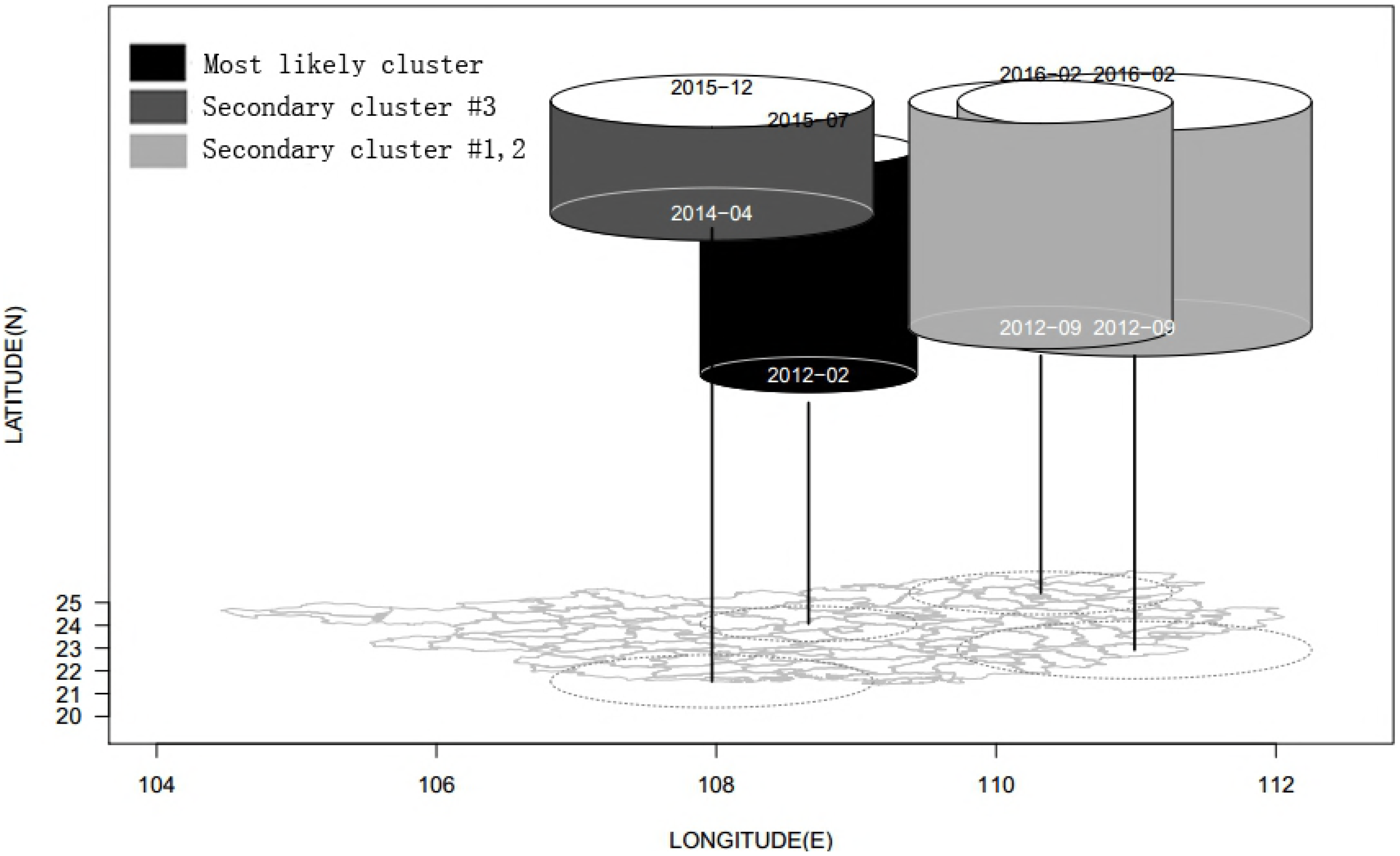
3D cluster window based on space-time scan in Guangxi, 2010-2016

### Spatial panel models

As shown in table 2, Moran’s I were greater than 0 from 2010 to 2016. It indicated the existence of high spatial dependency on the incidence of TB, thus we took the spatial autocorrelation into consideration. Three panel models with spatial lag and spatial error correlation were fitted (On effect model, fixed effects model and random effects model). The results are shown in Table 4. By comparing their rho (ρ), lambda (λ) as well as random effects chinking, we found that the random effects model is better than the other two. Duration of sunshine, per capita gross domestic product (PGDP), the recovery rate of TB and participation rate of new cooperative medical care insurance in rural areas had a significant negative association with the reported incidence of TB.

**Table 4.**
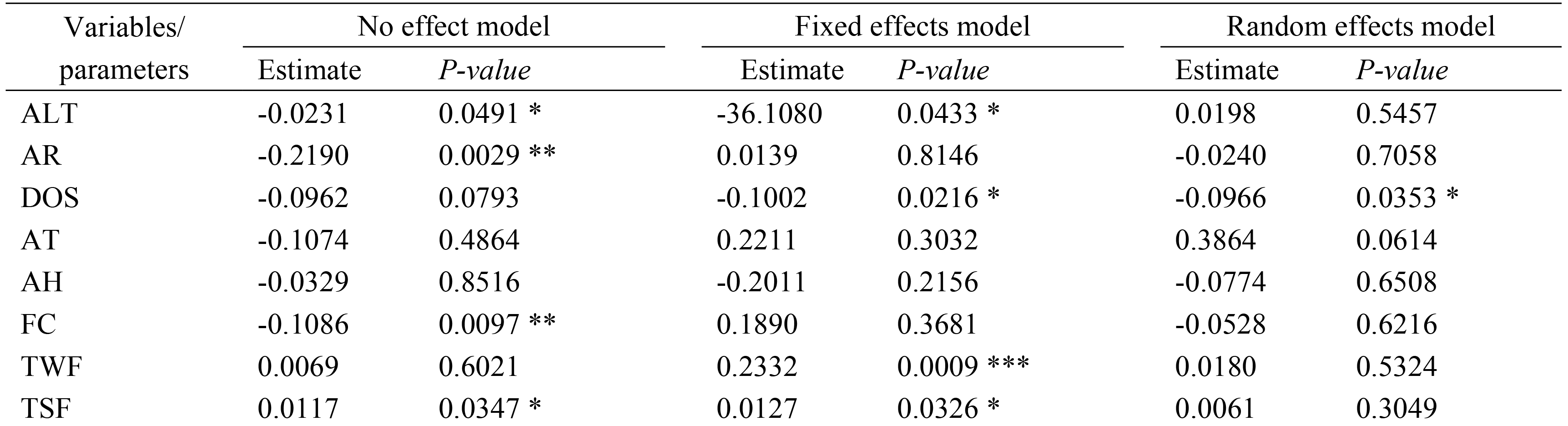
Results of the panel models with spatial lag and spatial error correlation

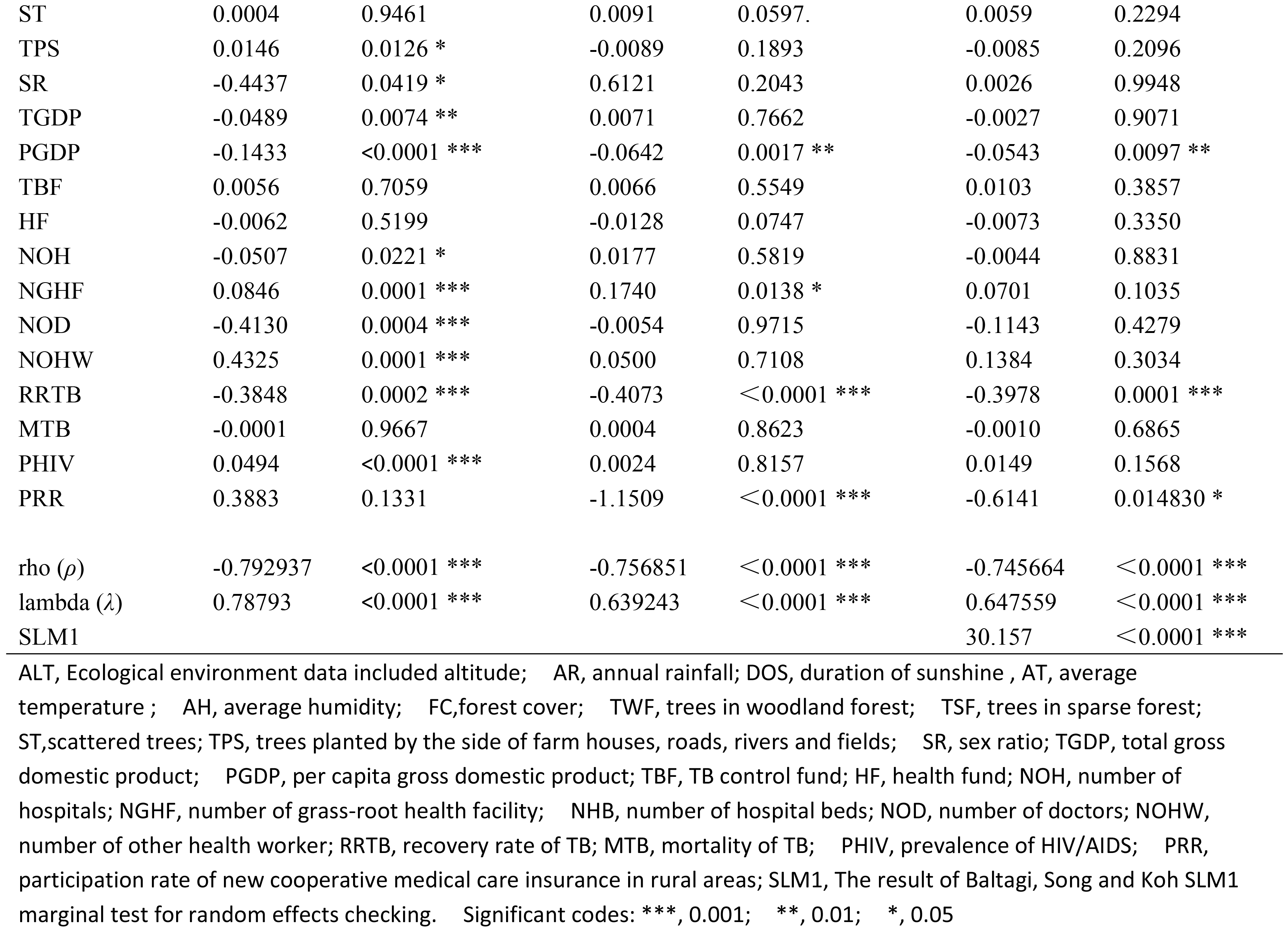

## Discussion

This study explored the spatiotemporal pattern of active TB transmission and its ecological causation using time series analysis, spatial autocorrelation analysis, spacetime scan statistics and spatial panel modeling. The number of active TB cases reported from NNDRS is getting closer to the number estimated by WHO. ^[1,18]^ Thus we feel the results of our study are close to the truth. To our knowledge, this the first study to conduct a spatial panel data analysis of active TB cases in Guangxi, China.

While the global incidence of TB has fallen by an average of 2% annually between 2010 and 2016, from our study, the number of active TB cases in Guangxi has decreased slowly. The annual incidence has remained at a high level (above 100 cases per 100,000 population), slightly lower than the global average of 133.

The times series of TB in Guangxi showed a seasonal trend over the seven-year period with peaks occurring in March and July. This result is similar to previous studies in Guangxi, Pakistan and Portugal.^[19–21]^ The number of TB cases in 2017 also exceeded 490,000 with an estimated incidence of 108.3 per 100,000 population. The peaks appeared in spring and summer, a pattern similar to India.^[22]^ Some researchers suggest that most TB patients who are infected with MTB in the winter then develop active TB during the next 3-6 months.^[23]^ Furthermore, the delay of TB diagnosis and reporting can also lead to the postponement of peaks.^[9]^

In this study, we applied *Moran’s I* statistics with EB adjustment to identify the global and local autocorrelation of TB cases at the county level. The statistics of the global *Moran’s I* ranged from 0.363 to 0.536 for all seven years, indicating that TB cases reported by the health department were clustered geographically at the county level with similar incidence rates. Furthermore, It shows high spatial dependency on the incidence of TB. The panel models with spatial lag and spatial error correlation were an excellent choice.^24^ The situation of TB clustering has existed for several years in Guangxi. Local patterns of TB cases display the actual location of high-high spots and low-low spots. From the local result, the most significant hot spots (high-high location) were mainly gathered in the central and western parts of Guangxi. Some neighboring districts (Du’an, Yizhou, Xincheng and Xingbin) always exhibited high-high pattern as the strongest hot spots for all seven years.

The research data has two dimensions (location-based variables and timeline). The space-time scan technique which enable researchers to detect the interaction of space and time has been widely used to find spatiotemporal clusters in cancers, including brain cancer,^[25]^ lung cancer,^[26]^ gastric cancer and breast cancer,^[27–28]^ and also infectious diseases such as hand-foot-mouth disease,^[29, 2]^ dengue fever and TB.^[30–31]^ A previous study of spatiotemporal clustering characteristics of TB in China showed that the most likely cluster included Guangxi had a higher TB burden and a higher risk of TB transmission between 2005 and 2011, suggesting that this area should be targeted for TB control and prevention. ^[7]^ By using the space-time scan statistic, we detected one most likely cluster and three secondary clusters in Guangxi between 2010 and 2016. These TB clusters were identified similar to that of the *Moran’s I* local autocorrelation statistics but introduced the time variable into the model. Xincheng county is the central point of the most likely cluster, located in the center of Guangxi with a high relative risk occurring between February 2013 and July 2015. The residents of nine neighboring counties in the most likely cluster had a 1.93 times higher risk of developing active TB compared to those living outside these counties. Compared with the high TB reported incidence population, the people living in the eastern areas had a lower risk of TB (RR=0.62, Table 3).

Spatial dependence might exist between the observations at each unit, especially in infectious disease monitoring data.^32^ It is the first time for this study to quantify the association between ecological environment factors and TB using spatial models on the basis of longitudinal data from 112 counties for the period 2010 to 2016. The study highlights the random effects of spatial panel models to determine the association between TB and ecological environment factors. The findings showed that the annual reported incidences of TB in Guangxi, China was significantly associated with Duration of sunshine, per capita gross domestic product (PGDP), the recovery rate of TB and participation rate of new cooperative medical care insurance in rural areas. A negative relationship between duration of sunshine and TB reported incidence was observed, which is consistent with the findings of other recent studies.^33,34^ It means that the less someone exposure in the sunshine, the higher risk of the TB he/she get. A mechanism for this relationship is that decreased vitamin D levels with consequent impaired host defense arising from reduced sunshine exposure.^35^ It is necessary to educate people to increase the outdoor exercise. The other significant negative variable is PGDP, which is one of the important indexes to measure the economic level. Since poverty was reported as one of the risk factors for TB in the USA,^[36]^ it would be a problem if the high TB incidence rate were associated with low-socioeconomic status in the middle Guangxi. The economies in eastern Guangxi (TB cold spots) which share a boundary with some counties of neighboring Guangdong province (a developed province in China) have developed rapidly. In this study, the lower recovery rate of TB and lower-level Medical insurance coverage are both lead to The long-term existence of the source of infection, which is similar to the results of recent studies.^[37,38]^ The situation of failed TB treatment and out-of-pocket payments with catastrophic consequences will hamper the efforts to end TB.

## Limitations

There are some limitations in this study. Firstly, the system was based on passive case findings, a method which is likely to miss some cases. Secondly, we conducted the space-time scan statistic based on a cylindrical scanning window. Since districts are irregular, this might have led to an erroneous judgment of the number of units in a cluster. Unfortunately, the *Flex-scan* for irregular spatial detecting cannot incorporate time. Thirdly, Some potential risk factors for causation such as environmental pollution and socio-economic factors, which may be associated with the TB clustering, were not included in the spatial model. It due to the lacking of monitoring stations at the county level.^[39–41]^

## Conclusion

The situation of TB in Guangxi remains serious. This study detected a spatial and temporal pattern of TB transmission in Guangxi Zhuang autonomous region using spatiotemporal statistics. The main cluster was located in the central part of Guangxi. Spatial panel modeling with random effects identified that duration of sunshine, per capita gross domestic product (PGDP), the recovery rate of TB and participation rate of new cooperative medical care insurance in rural areas had a significant negative association with the reported incidence of TB. Promoting the productivity, improving TB treatment pathway and strengthening environmental protective measures (increasing sunshine exposure) are the key wards to TB control.

## Declarations

### Ethical review

This research was approved by the Institutional Review Board of the Guangxi (GW-2017-0001). Ethics committees approve this consent procedure. All participants provided their written informed consent to participate in this study.

### Consent for publication

Not applicable.

### Availability of data and material

We collected the data from NNDRS, those reported by the General Hospital, TB designated hospitals, CDC and specialist hospitals and verified by NTP. Ecological data were obtained from the China Meteorological Data Sharing Service System, statistical yearbook and health resource database.

### Authors’ Contributions

Conceived and designed the experiments: D.L., Z.C., V.C. Collected data: Z.C., D.L. J.Z. Analyzed the data: Z.C., D.L., V.C., M.L. Contributed reagents/ materials/ analysis tools: Z.C., J.O. J.Z. Wrote the paper: Z.C., D.L. All authors read and approved the final manuscript, and accepted the accountability for all aspects of the work.

## Acknowledgments

Advice on statistical analysis was provided by Edward McNeil from the Epidemiology Unit, Faculty of Medicine, Prince of Songkla University, Songkhla, Thailand. We would also like to thank staff from all centers for disease prevention and control in Guangxi (city level and county level) for all their patience and support. This study was a part of the Ph.D. thesis of the first author to fulfill the requirement for the TB/MDR-TB research training program at Epidemiology Unit, Prince of Songkla University under the support of Fogarty International Center, National Institutes of Health (D43TW009522).

